# Synthesis, Characterization, and Nanodisc formation of Non-ionic Polymers

**DOI:** 10.1101/2021.01.11.426267

**Authors:** Thirupathi Ravula, Ayyalusamy Ramamoorthy

**Author notes:** Ayyalusamy Ramamoorthy: Author info 1.

## Abstract

Despite lipid-nanodiscs are increasingly used in the structural studies of membrane proteins, drug delivery and other applications, the interaction between the nanodisc-belt and the protein to be reconstituted is a major limitation. To overcome this limitation and to further broaden the scope of nanodiscs, a family of non-ionic amphiphilic polymers synthesized by hydrophobic functionalization of fructo-oligosaccharides/inulin is reported. We show the stability of lipid-nanodiscs formed by these polymers against pH and divalent metal ions, and their magnetic-alignment properties. The reported results also demonstrate that the non-ionic polymers extract membrane proteins with unprecedented efficiency.

---

Recent studies have successfully demonstrated the use of synthetic polymers to directly extract membrane proteins and reconstitute them in near-native lipid bilayer nanodiscs without using a detergent.^1-7^ However, the presence of charge on the currently known polymers drastically limits their applications.^8, 9^ The high charge density of the polymer interferes with the purification by ion-exchange chromatography and also detrimental to study oppositely charged membrane proteins due to charge-charge interactions.^8, 9^ Therefore, there is considerable interest in developing non-ionic polymers to expand the applications of polymer-based nanodiscs technology for structural biology, drug delivery, biosensors and other technological applications. Here, we show that naturally-extracted oligosaccharides can be used as the polymer backbone which upon hydrophobic functionalization form lipid nanodiscs.

Fructo-oligosaccharides (FOS) or fructans are the natural biopolymers found in chicory root, garlic, onion and other fruits.^10^ The degree of polymerization of FOS ranges from 2 to about 60;^11^ the high molecular weight FOS are called inulin. In this study, inulin obtained from chicory was used as the starting material for hydrophobic functionalization. After MALDI-MS (matrix-assisted laser desorption ionization mass spectrometry) characterization, that revealed the average degree of polymerization (DP) to be ∼14, inulin was functionalized with pentyl bromide in the presence of sodium hydride (Figure 1A). The resulting polymer was characterized by NMR and MALDI mass spectrometry. The degree of substitution (DS), which is the average number of functional groups attached per one fructose monomer, was estimated using ^1^H NMR by integrating the peak from H1-Glc (at 5.4 ppm) which was then used as the reference to quantify the extent of functionalization by using the peak at 0.9 ppm from the terminal methyl group of the pentyl group (See Figure S1). The resulting polymer was further characterized by MALDI-MS and solution and solid-state NMR experiments. MALDI-MS spectra showed an average molecular weight (MW) of 2.7 kDa. The ^13^C CP-MAS and 2D ^1^H-^13^C HSQC spectra revealed the presence of pentyl groups showing the successful functionalization.

**Figure 1.**
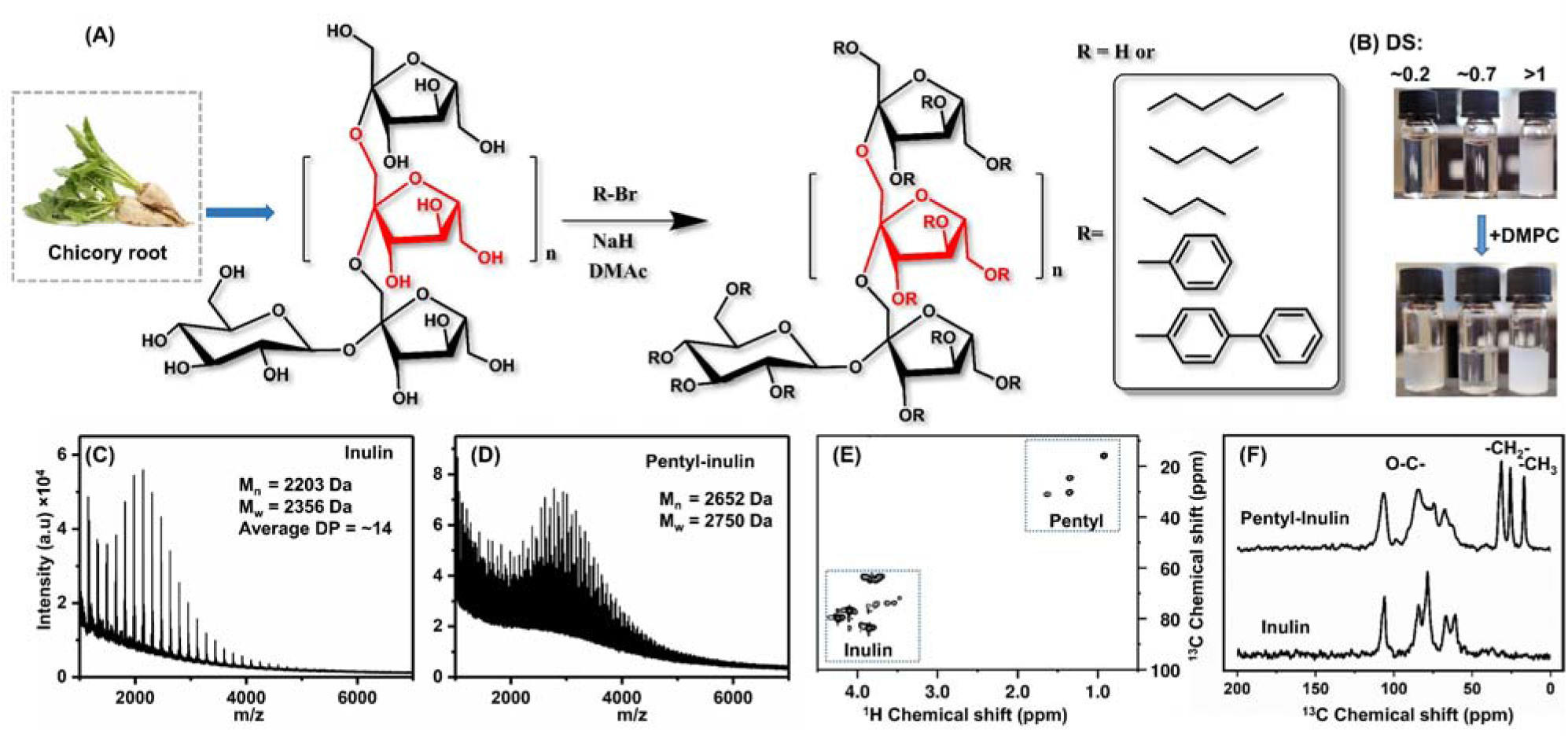
Design, synthesis and characterization of non-ionic fructo-oligosaccharide polymers. (A) A synthetic scheme showing the reaction to hydrophobic modification of inulin extracted from chicory. (B) The solubility of pentyl-inulin derivatives with a varying degree of substitution (DS=∼0.2, ∼0.7, and >1) in water (5 mg/ml). (C, D) MALDI-MS spectra of inulin (C) and pentyl-inulin (D) showing the degree of polymerization. E) 2D ^1^H-^13^C-HSQC solution NMR spectrum of pentyl-inulin and (F) ^13^C CP-MAS spectra of Inulin (bottom) and pentyl inulin (top) acquired under 12 kHz spinning showing the presence of peaks from the pentyl functional groups and thus confirming the extent of functionalization of inulin.

One of the main constraints in the design of an amphiphilic polymer to form nanodiscs is that there should be an optimal ratio of hydrophilic to hydrophobic moieties.^12^ In the case of inulin, this ratio is represented by DS which is the average number of hydrophobic units per fructose monomer. The solubility of pentyl functionalized inulin in water was found to depend on DS as shown in Figure 1B. The resulting polymers with DS<1 was found to be soluble in water, whereas those with DS>1 were found to be insoluble and therefore not considered for further studies.

Inulin polymers with DS values 0.2 and 0.7 were tested for their ability to dissolve lipid aggregates by static light scattering (SLS) experiments. Pentyl-inulin (with DS 0.2 and 0.7) was added to DMPC (1,2-dimyristoyl-sn-glycero-3-phosphocholine) liposomes and SLS profiles were measured as a function of time. DMPC liposomes showed a high light scattering intensity due to their large size in the absence of polymer; which upon the addition of a polymer (with DS 0.7, at polymer to lipid ratio of 1:1 w/w) exhibited a steep decrease in the light scattering intensity suggesting the polymer-induced formation of smaller particles. The resulting solution became completely transparent (Figure 1B). The extent of decrease in static light scattering intensity was found to depend on the lipid:polymer ratio. It should be noted that when a polymer with a low degree of substitution was used, the light scattering intensity showed only a partial solubilization of DMPC liposomes (Figure 2B). These results indicate that polymers with a DS value of 0.7 is optimal for the solubilization of DMPC liposomes. Using the DS values within 0.7-1 range as the optimal amount of functionalization, we then functionalized with various alkyl (butyl and hexyl) and aromatic (benzyl and biphenyl) moieties. The resulting polymers were then characterized using MALDI-MS, ^13^C CP-MAS solid-state and 2D ^1^H-^13^C HSQC solution NMR experiments (Figures S2-S10).

**Figure 2.**
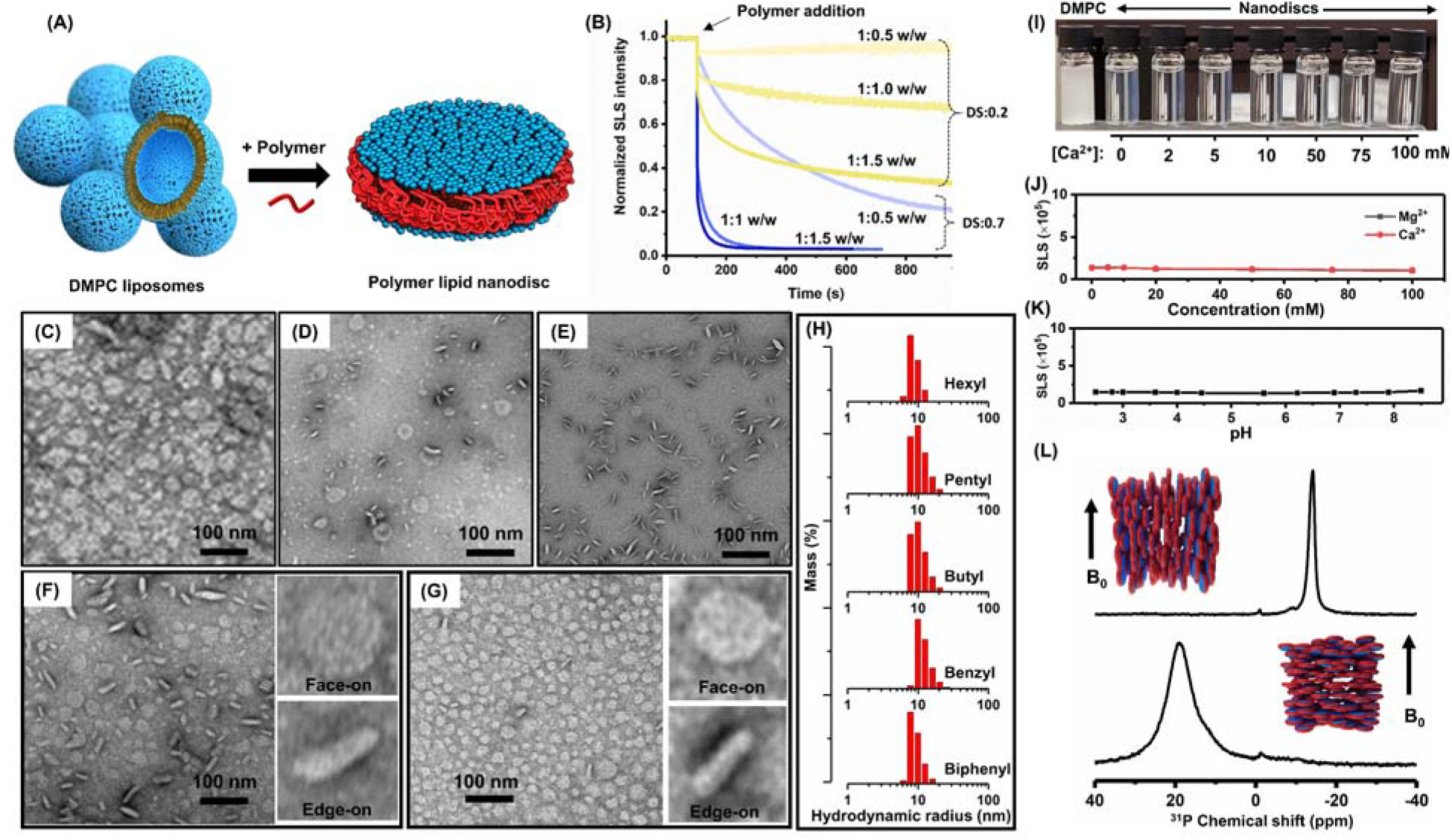
Solubilization of liposomes to form nanodiscs by non-ionic polymer. (A) Schematic representation of the formation of a polymer-based lipid-nanodisc from liposomes. (B) Static light scattering experimental profiles showing the solubilization of DMPC liposomes at various polymer concentrations (indicated as DMPC:polymer w/w ratio); data for two different degrees of substitution (DS = 0.2 and 0.7) of polymers are indicated. The arrow mark in (B) indicates the time at which the polymer was added. (C-G) TEM images of nanodiscs prepared from 1:1 w/w lipid:polymer: (C) pentyl-inulin:DMPC, (D) Hexyl-inulin:DMPC, (E) Benzyl-inulin:DMPC, (F) Pentyl-inulin:DMPC-DMPG (7:3), and (G) Hexyl-inulin:DMPC-DMPG (7:3). Insets in (F and G) show nanodiscs face-on (top) and edge-on (bottom) sides of the nanodisc. Scale bar represents 100 nm. (H) DLS profiles of nanodiscs prepared from 1:1 w/w lipid (DMPC:DMPG (7:3)) to polymer ratio for the indicated polymers showing the hydrodynamic radius of the nanodiscs. (I) A photograph of nanodiscs (prepared by pentyl-inulin:DMPC (1:1 w/w)) solution in the presence of various concentrations of Ca^2+^. Static light scattering intensity of the nanodiscs in the presence of various concentrations of Ca^2+^ or Mg^2+^ (J), and for varying pH values (K). (L) ^31^P NMR spectra of magnetically-aligned nanodiscs: (top) with the bilayer-normal perpendicular to the external magnetic field and (bottom) in the presence of 2 mM YbCl_3_ flipped to have the bilayer-normal parallel to the external magnetic field. Inset showing the schematic representation of the orientation of nanodiscs with respect to the applied magnetic field direction.

We then characterized the ability of the polymers to solubilize the liposomes to form nanodiscs using TEM and DLS. DLS profiles of DMPC:DMPG (1,2-dimyristoyl-sn-glycero-3-phospho-(1’-rac-glycerol)) (7:3 molar ratio) nanodiscs formed by various polymers (Figure 2H) showed a hydrodynamic radius of ∼10 nm. The TEM images showed circular discs with face-on edge-on for DMPC (Figure 2C-E) as well as for DMPC-DMPG (Figure 2F and 2E). These results demonstrate the: (a) hydrophobic functionalization of inulin polymers enables the formation of nanodiscs, (b) hydrophobic moiety can be changed to various functional groups such as alkyl chains or aromatic moieties, and (c) nanodiscs formation by the reported polymers with different lipid composition.

One of the major limitations of many of the currently used nanodiscs is their poor stability against pH and divalent metal ions.^1, 13^ Therefore, we tested the stability of the non-ionic inulin-based polymer nanodiscs in the presence of divalent metal ions and also at various pH values. The SLS profiles show no difference in intensity as a function of pH and at various concentrations of Ca^2+^ and Mg^2+^ (Figure 2I-2K). These results indicate that the inulin based nanodiscs are stable under a pH range of 2.5-8.5 and also up to 100 mM divalent metal ion concentrations.

An unique property of large-size (>20 nm in diameter) nanodiscs (called as macro-nanodiscs) is their ability to align in the presence of an external magnetic field. This is a much needed property to study membrane proteins using solid-state NMR techniques;^14-16^ and also useful for solution NMR applications via the measurement of residual dipolar couplings (RDCs)^17-20^ from water-soluble molecules by utilizing nanodiscs as the alignment medium.^21^ To examine the magnetic-alignment properties of the non-ionic inulin based nanodiscs, static solid-state NMR experiments were performed. ^31^P NMR spectra of nanodiscs made from pentyl-inulin with DMPC (1:1 w/w) showed a peak at ∼-14±0.5 ppm suggesting that the nanodiscs are aligned with the lipid-bilayer-normal perpendicular to the magnetic field direction (Figure 2L). In addition, the ability to flip the alignment direction by 90° is also demonstrated by adding 2 mM YbCl_3_ to the sample, which showed a ^31^P NMR peak at ∼19±3 ppm suggesting that the bilayer-normal is oriented parallel to the applied magnetic field axis. These NMR results demonstrate that the non-ionic polymers can be used for structural studies of membrane proteins by solid-state NMR spectroscopy and RDCs based solution NMR studies.^14, 16, 21-23^

One of the applications of synthetic nanodisc-forming polymers is their ability to directly extract membrane proteins from the native environment without the use of a detergent.^1^ To demonstrate the feasibility of using the non-ionic inulin based polymers for this application, *E*.*coli* membranes were incubated with each of the reported inulin based polymers and the concentration of solubilized membrane proteins were estimated using bicinchoninic acid (BCA) assay (Figure S11).^24^ Figure 3 shows the solubilization efficiency of the inulin based and other nanodisc-forming polymers with that of a commonly used detergent DDM (dodecyl maltose). All of our nonionic inulin-based polymers showed an efficacy of >80% as compared to DDM’s 100% efficiency. Pentyl-inulin showed the highest efficiency when compared to other polymers. Whereas the currently used charged polymers showed only <50% efficiency compared to DDM detergent. The higher efficiencies of the inulin polymers reported in this study can be attributed to the absence of charge in them which enables them to solubilize both anionic and cationic proteins. These results indicate that the polymers developed in this study have a great potential for detergent-free extraction of a variety of membrane proteins. In addition, it is worth mentioning that the absence of a chromophore in the alkyl functionalized polymers synthesized in this study makes them excellent chemical tools as they do not interfere with the absorption/emission properties of the reconstituted membrane proteins in biophysical characterization.

**Figure 3.**
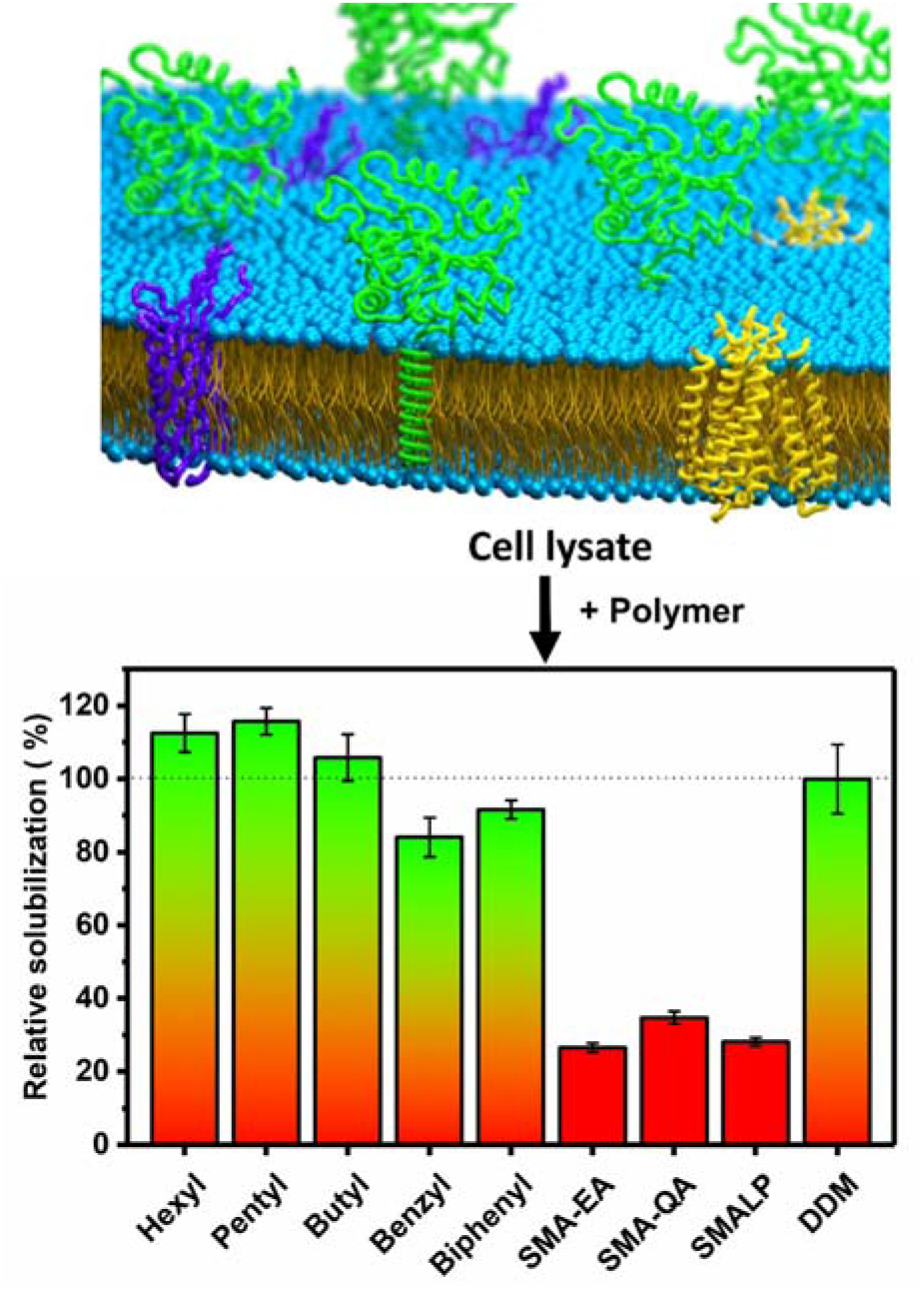
Membrane solubilization by non-ionic polymers. Schematic representation and relative solubilization by various polymers compared using the BCA assay (Figure S10).

In conclusion we have successfully developed the first non-ionic polymers that are shown to form nanodiscs. The reported results demonstrate that: (a) the efficacy of membrane solubilization depends on the degree of substitution in the polymer, (b) various hydrophobic moieties can be used to functionalize inulin, (c) the inulin based nanodiscs are stable under different pH values and various concentrations of divalent metal ions, (d) the inulin based nanodiscs magnetically-align in the presence of an external magnetic field which can be utilized in the structural studies of membrane proteins by NMR, and (e) the new polymers can be used for detergent-free extraction and purification of membrane proteins irrespective of their charge. Thus, the newly developed non-ionic polymers would be useful to expand the applications of the nanodisc technology.^25-31^ Further, the natural source of the inulin-based polymer backbone makes them a great candidate for further developments of biocompatible polymers and will also find many applications in drug delivery and membrane protein structural biology.

## Acknowledgements

This study was supported by NIH (R35GM139573 to A.R.). Authors declare that they have included the inulin based polymers in a recently filed US patent.

