## Supporting information for "Synthesis, Characterization, and Nanodisc formation of Non-ionic Polymers"

### **Materials and Methods**

Inulin from chicory, anhydrous dimethylacetamide (DMAc), potassium phosphate, hydrochloric acid (HCl), and sodium hydroxide (NaOH), diethyl ether (ether), calcium chloride (CaCl_2_), magnesium chloride (MgCl_2_), sodium chloride (NaCl), phosphoric acid (H_3_PO_4_), potassium phosphate, sodium hydride, pentylbromide, butylbromide, Benzyl bromide and hexylbromide were purchased from Sigma-Aldrich® (St. Louis, MO, USA). 1,2-dimyristoyl-sn-glycero-3-phosphocholine (DMPC) and 1,2-dimyristoyl-sn-glycero-3-phospho-(1'-rac-glycerol) (DMPG) were purchased from Avanti Lipids Polar, Inc®. Hydrochloric acid (HCl), sodium hydroxide (NaOH), 4-Bromomethylbiphenyl and Pierce™ BCA Protein Assay Kit were purchased from Thermo Fisher Scientific® (Waltham, MA, USA). SMA-3000 was a kind gift from Polyscope (Netherlands).

**Hydrophobic modification of inulin:** 1 g of inulin (5.5 mmol of fructose) was added to 30 ml of dimethylacetamide (DMAc), followed by heating the reaction mixture to 50 °C under stirring until complete solubilization of inulin. Then, the reaction mixture was cooled to room temperature and sodium hydride (2 eq, 442 mg, 60% NaH in mineral oil) was added slowly in three portions over 1 hr duration. The resulting mixture was heated to 50 °C and stirred for 1 hr. Then, the reaction mixture was allowed to cool to room temperature, followed by the addition of 1.5 eq (8.3 mmol) of alkyl bromide (1.04 ml of pentylbromide; 0.9 ml of butyl bromide; 1.16 ml of hexyl bromide; 0.98 ml of benzyl bromide; 2g of 4-Bromomethylbiphenyl; The degree of substitution can be controlled by the amount of the alkyl bromide). Then the reaction mixture was stirred at room temperature. The reaction progress was monitored by taking a 100 µl of aliquot of reaction mixture, precipitated with ether and ethanol (50% v/v). The sample was allowed to dry under a stream of nitrogen gas followed by drying under high vacuum. The resulting powder was characterized by ^1^H NMR by dissolving it in D_2_O. The progress of the reaction was monitored over two days and the reaction was stopped when the required degree of functionalization was achieved. After reaching the required degree of functionalization, the reaction was quenched by the addition of ice-cold ethanol, followed by the addition of diethyl ether. The resulting precipitate was separated by slow centrifugation (3000 rpm for 5 min). The precipitate was washed three times with ether and followed by washing with ethanol three times to remove any low molecular weight species.^[1]^ The resulting precipitate was dried under high vacuum (yield 70-80%). The resulting polymer was dissolved in ice cold water and the pH of the solution was adjusted by the addition of 0.1 M HCl and then lyophilized to obtain a white powder. The polymer was characterized using MALDI-MS, 1D ^1^H and 2D ^13^C-^1^H HSQC and ^13^C CP-MAS NMR experiments.

**Preparation of nanodiscs:** Polymer stock solutions were prepared by dissolving 10 mg of polymer in 1 mL of 20 mM Tris 50 mM NaCl (pH 7.4). DMPC stock solutions were prepared by the addition of 10 mg of DMPC powder in 1 mL of 20 mM Tris 50 mM NaCl (pH 7.4) and subjected to three freeze-thaw cycles to get a milky white solution. Nanodiscs were prepared by the addition of polymer to DMPC solution with weight ratios corresponding to 1:1 w/w of DMPC and polymer, and then incubated overnight at 4 °C.

**Matrix-assisted laser desorption/ionization mass spectrometry (MALDI-MS):** MALDI-MS was performed using Bruker spectrometer. Sample matrix 2,5-dihydroxybenzoic acid (DHB) was dissolved in Milliq water to give final concentration of 10 mg/mL. Inulin and its functionalized derivatives were dissolved in water to prepare a 1 mg/mL solution. The samples were added to the matrix solution to give 1:5 v/v. 1 µL of sample + matrix mixture was placed on a stainless steel target plate and air dried. External calibration was done using cyt C samples. MALDI-MS spectra were acquired using Bruker AutoFlex Speed operating at 20 keV accelerating potential. Spectra were acquired on positive ion mode, data were collected by obtaining 500-1000 shots until a reasonable signal-to-noise ratio was obtained by rastering on the whole sample to get reasonable signal to noise ratios. Molecular weight distributions were obtained using PolyTools™ software (Bruker Daltonik GmbH).

#### **Dynamic Light Scattering (DLS):** All DLS experiments were performed using Wyatt Technology® DynaPro® NanoStar® with a 1 μL quartz MicroCuvette.

#### **Transmission Electron Microscopy (TEM):** The TEM micrographs were obtained using a Technai® T-20® machine (FEI®, Netherlands) with 60 keV operating voltage. A dilute solution was dropped on carbon-coated copper grids and dried overnight at room temperature in a desiccator before use in the experiments.

**^1^H-NMR spectroscopy:** 25 mg of dried polymer was dissolved in 600 µl of D_2_O. NMR spectra were acquired using a 500 MHz Bruker NMR spectrometer.

**CP-MAS solid-state NMR experiments:** Carbon-13 CP-MAS spectra were acquired using a Bruker 400 MHz solid-state NMR spectrometer under 12 kHz MAS using a 2.5 mm triple-resonance MAS probe operating at 400.111 MHz and 100.617 MHz for ^1^H and ^13^C nuclei, respectively. Samples were packed into 2.5 mm zirconium oxide rotors. ^13^C CP-MAS spectra were obtained using the following experimental parameters: 2.5 µs 90° pulse, 2 ms CP contact time, 30 ms acquisition time, 10,000 scans, 5 s recycle delay and a 58 kHz radio-frequency decoupling of protons during the acquisition of ^13^C magnetization. ^13^C chemical shifts were calibrated to the chemical shift of CH_2_ resonance from a adamantane powder sample to 28.5 ppm.

**^13^C-^1^H Heteronuclear Single-Quantum Coherence (HSQC):** NMR samples were prepared by dissolving a 25 mg of polymer in D_2_O. Two-dimensional ^13^C/^1^H HSQC spectra were acquired on a Bruker 500 MHz NMR spectrometer equipped with a 5 mm triple-resonance TXI room temperature probe. Spectra were acquired using the following parameters: number of scans 32, 2048 t2*256 t1 points, spectral width 16 and 200 ppm in the ^1^H and ^13^C dimensions respectively, and 1 s recycle delay. The resulting spectra were processed using Topspin (from Bruker NMR) with zero-filling up to 2048*2048 points.

**^31^P NMR experiments:** ^31^P NMR experiments were acquired on a Bruker NMR spectrometer operating at a resonance frequency of 400.11 MHz for proton and 161.97 MHz for ^31^P nuclei. A 5 mm double-resonance MAS NMR probe was used under static condition. ^31^P NMR spectra were acquired using a 5 μs 90° pulse followed by a 25 kHz TPPM (two pulse phase modulation) proton decoupling. 512 scans were acquired for each sample with a relaxation/recycle delay of 2.0 s.

**Static Light Scattering (SLS) solubilization experiments**: The solubilization of DMPC MLVs was monitored by the intensity of scattered light at 90° angle. 3.5 mg/mL DMPC stock solutions were placed in a 4 mL cuvette (1 cm optical path) under continuous stirring at 20°C. Then the solution was equilibrated for 5 min before the addition of polymers. Excitation wavelength was set at 400 nm while the emission wavelength was set at 404 nm and the slit was set to 2 nm. All SLS experimental measurements were carried out using a FluoroMax-4® Spectrofluorometer from Horiba Scientific®.

**SLS metal ion titrations:** The stability of nanodiscs was tested by titrating a 3.5 mg/mL solution of Pentyl-inulin:DMPC (1:1 w:w) nanodiscs in 10 mM Tris buffer at pH = 7.4 with 2 M CaCl_2_ or 2 M MgCl_2_.

**pH stability measurement using SLS experiments:** To examine the pH stability of polymer-based lipid-nanodiscs, a solution made by 3.5 mg/mL Pentyl-inulin:DMPC (1:1 w:w) nanodiscs was titrated both with 1 M HCl and 1 M NaOH.

**Solubilization of native E. coli membranes**: *E. coli* BL21(DE3) cell pellets were re-suspended in a 10-fold volume of ice-cold buffer (20 mM Tris, 1 mM EDTA, 50 mM NaCl, pH 7.4) and subjected to ultrasonication. The cell lysate was subjected to centrifugation for 1 h at 4 °C and 20,000 g, and washed three times with buffer to remove the soluble proteins. Membrane pellets were re-suspended in 20 mM Tris buffer, 50 mM NaCl, pH 7.4 to a final concentration of 20 mg/mL (wet pellet weight). Solubilization of the pellet was carried out by treating 100 µL of cell lysate with 50 µL of polymer solution (20 mg/mL in 20 mM Tris, 50 mM NaCl, pH 7.4) and 50 µL of buffer (20 mM Tris, 50 mM NaCl, pH 7.4). DDM solubilization was performed by using 20 mM stock solution. The samples were incubated overnight at room temperature while shaking. The resulting solutions were centrifuged at 11,000 g for 30 min. The supernatant was separated and analyzed using Pierce™ BCA Protein Assay Kit using the recommended protocol by the supplier.

**Schematics:** Chemical structures in Figure 1A are drawn using ChemDraw® (PerkinElmer Informatics). The image of Chicory root shown in Figure 1A was taken from <https://sc02.alicdn.com/kf/HTB1iTdToDnI8KJjSszgq6A8ApXau.jpg>. Schematics used in Figures 2A and 3 were prepared using 3D modelling software Blender (<https://www.blender.org/>).

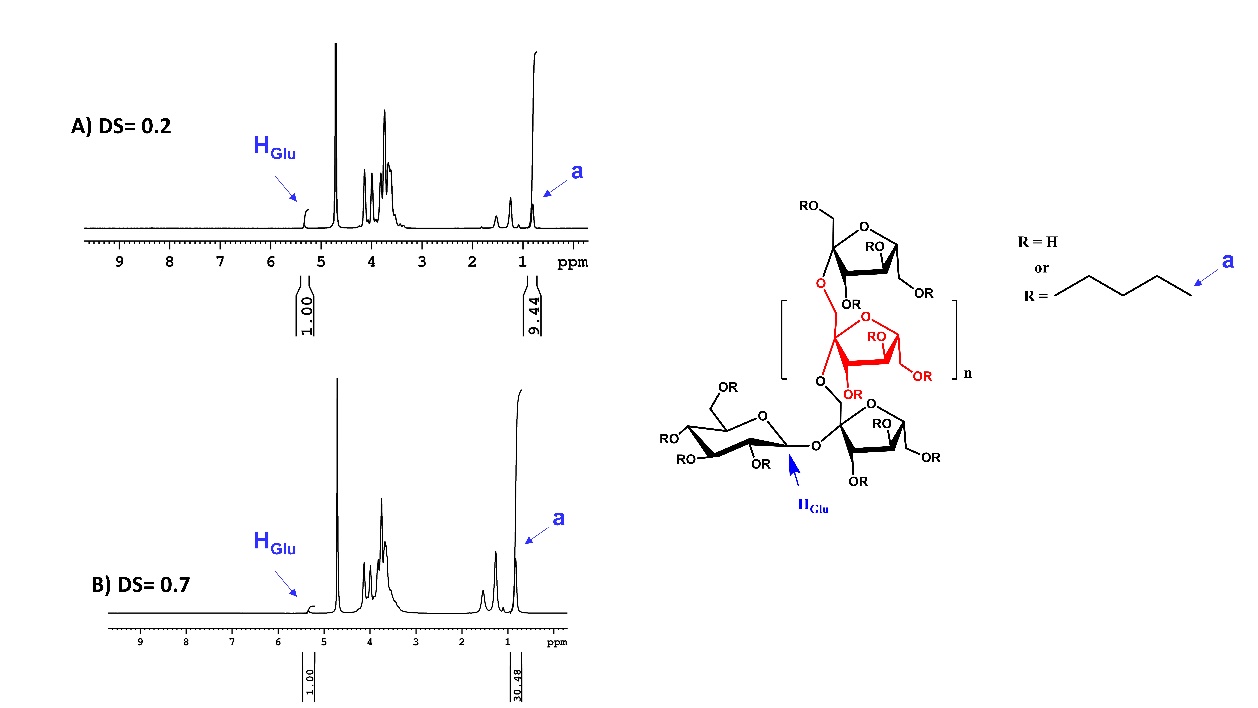

**Figure S1.** ^1^H NMR of pentyl-inulin polymer with different degree of substitution (DS of 0.2 and 0.7). The assignment of H_Glu_ and the terminal methyl groups (marked as ‘a’) that are used to estimate the extent of functionalization are shown. This method has previously been used to estimate the degree of substitution of inulin derivatives in the literature.^[2]^

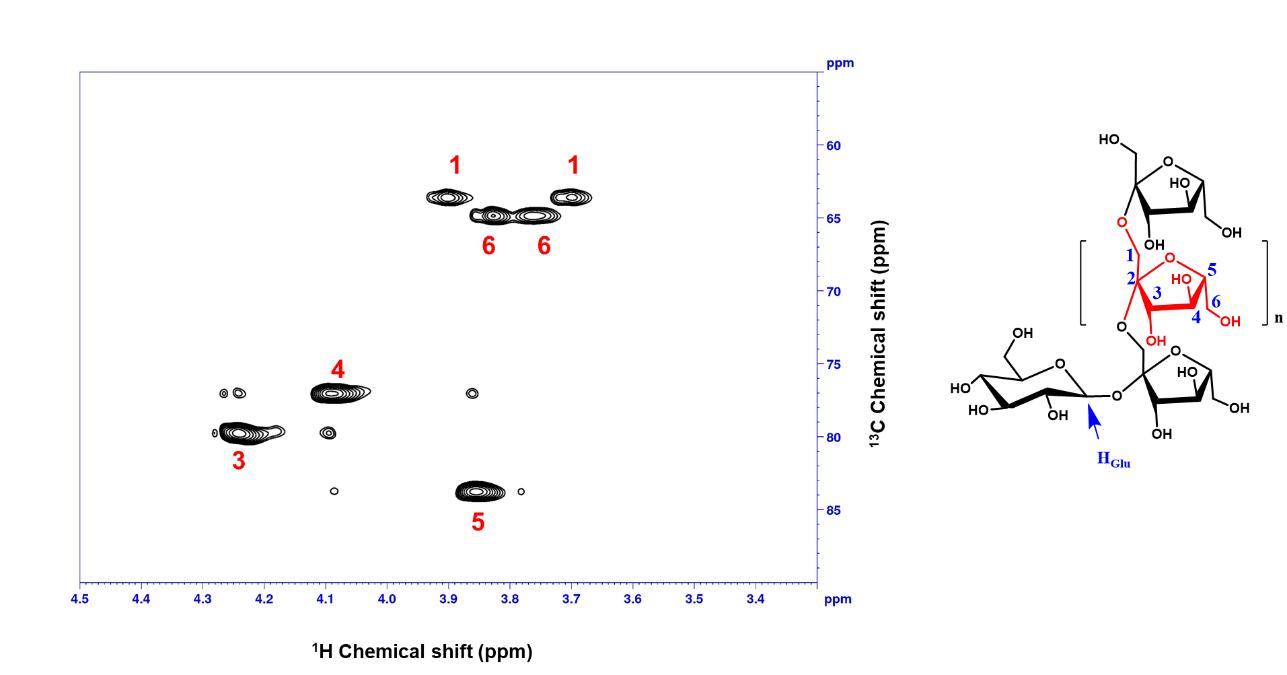

**Figure S2.** 2D ^13^C-^1^H HSQC NMR spectrum of inulin (25 mg/ml) dissolved in D_2_O shown along with resonance assignment.

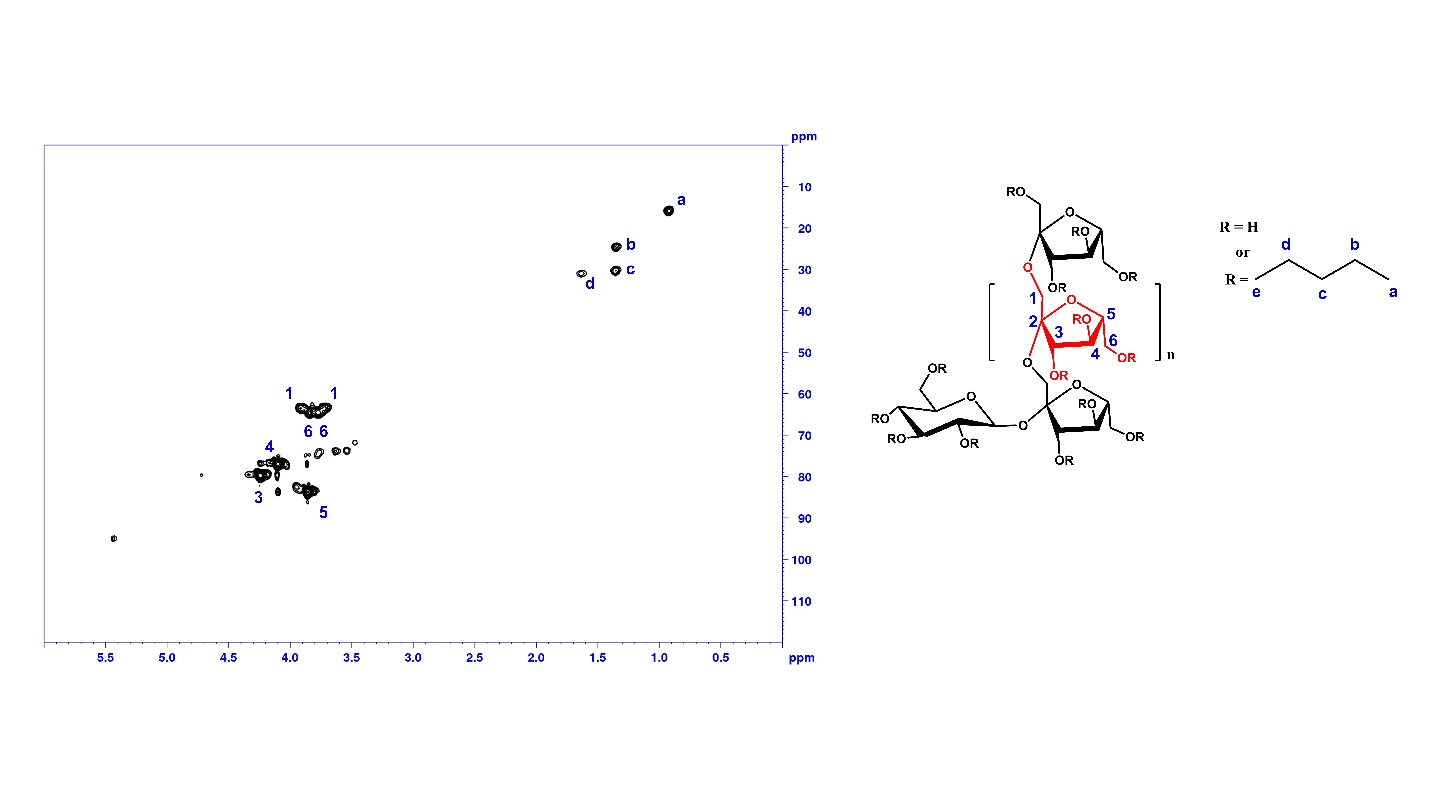

**Figure S3.** 2D ^13^C-^1^H HSQC NMR spectrum of pentyl-inulin (25 mg/ml) dissolved in D_2_O shown along with resonance assignment. Peaks from methyl ^1^H-^13^C (labeled as ‘e’) from the R groups are not assigned due to spectral overlap.

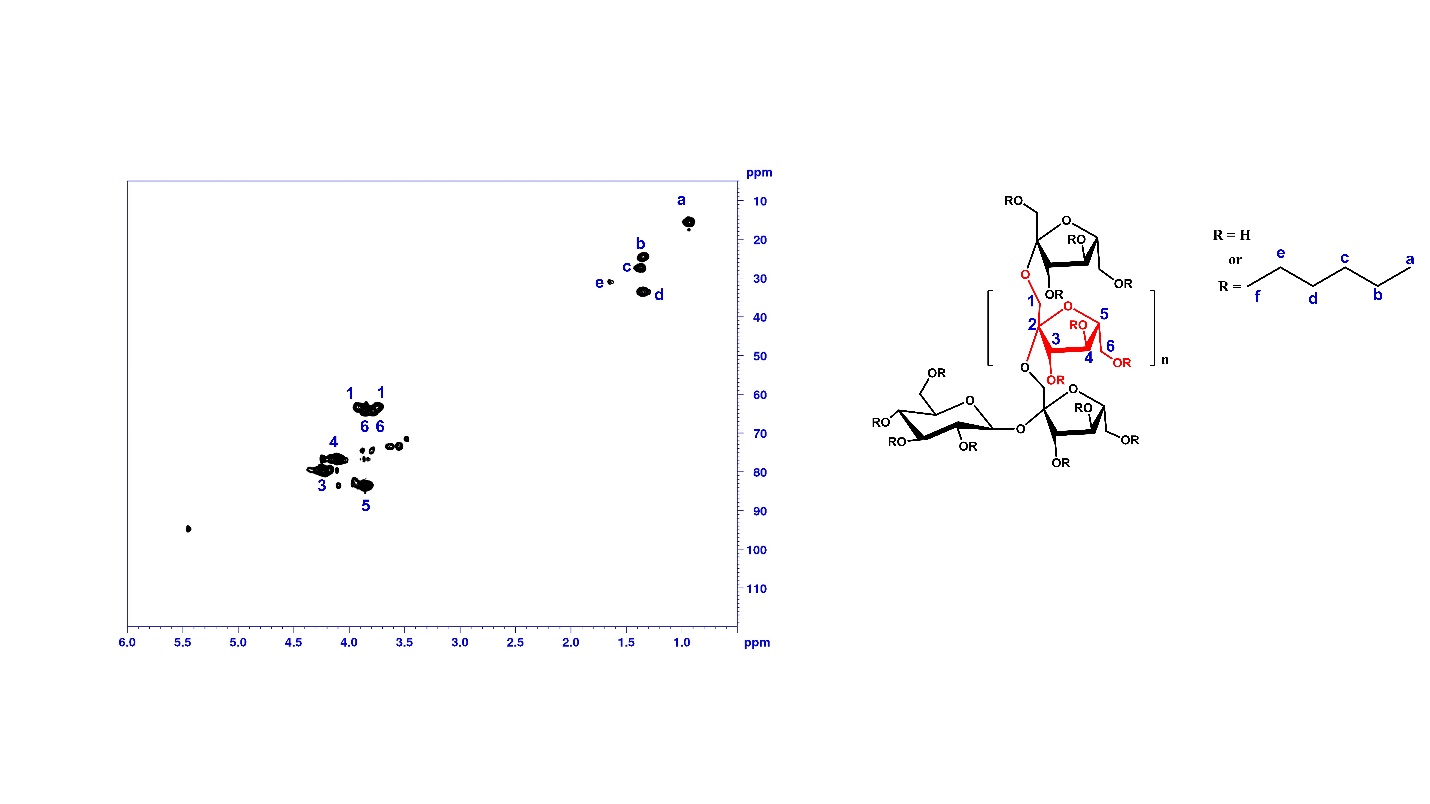

**Figure S4.** 2D ^13^C-^1^H HSQC NMR spectrum of hexyl-inulin (25 mg/ml) dissolved in D_2_O shown along with resonance assignment. Peaks from methyl ^1^H-^13^C (labeled as ‘f’) from the R groups are not assigned due to spectral overlap.

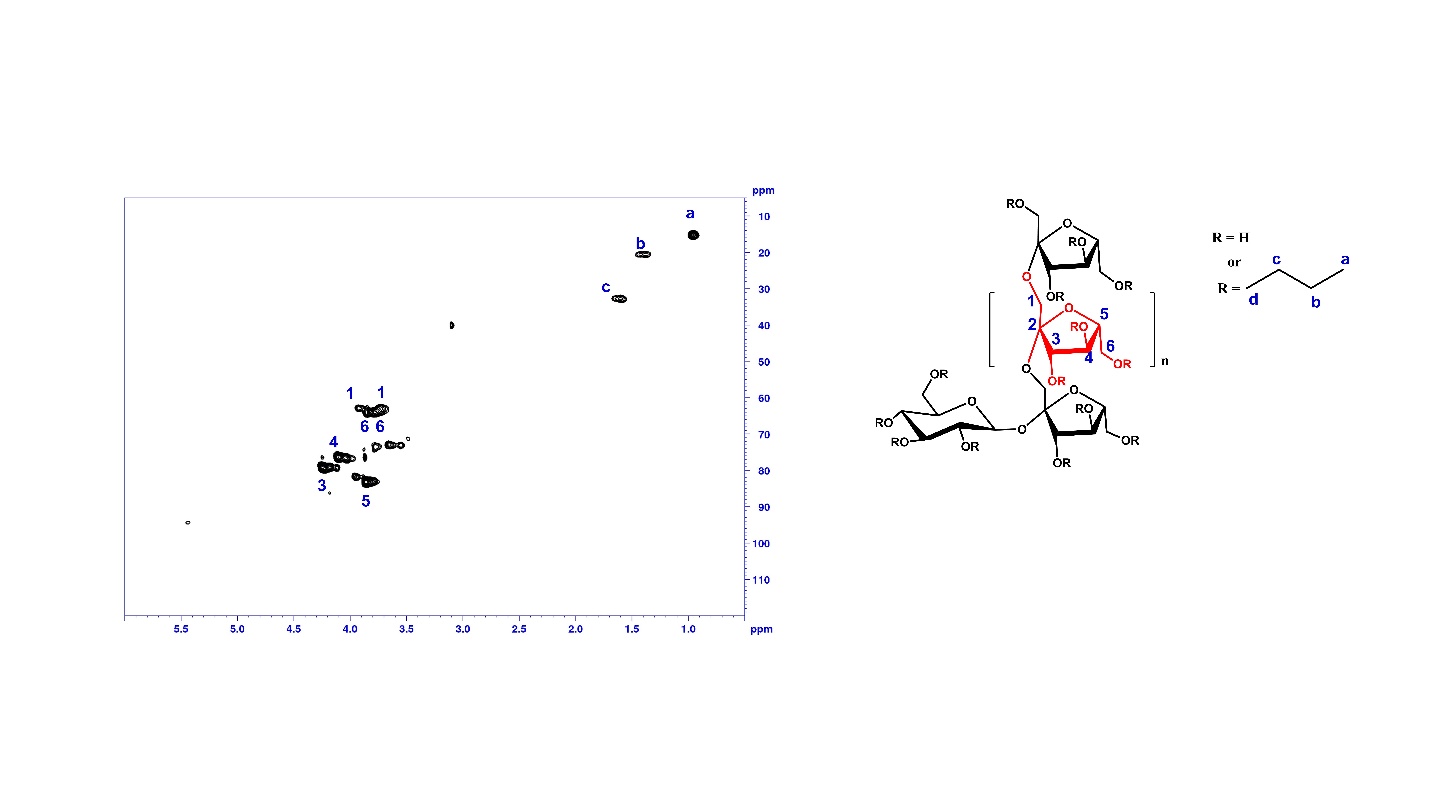

**Figure S5.** 2D ^13^C-^1^H HSQC NMR spectrum of butyl-inulin (25 mg/ml) dissolved in D_2_O shown along with the resonance assignment. Peaks from methyl ^1^H-^13^C (labeled as ‘d’) from the R groups are not assigned due to spectral overlap.

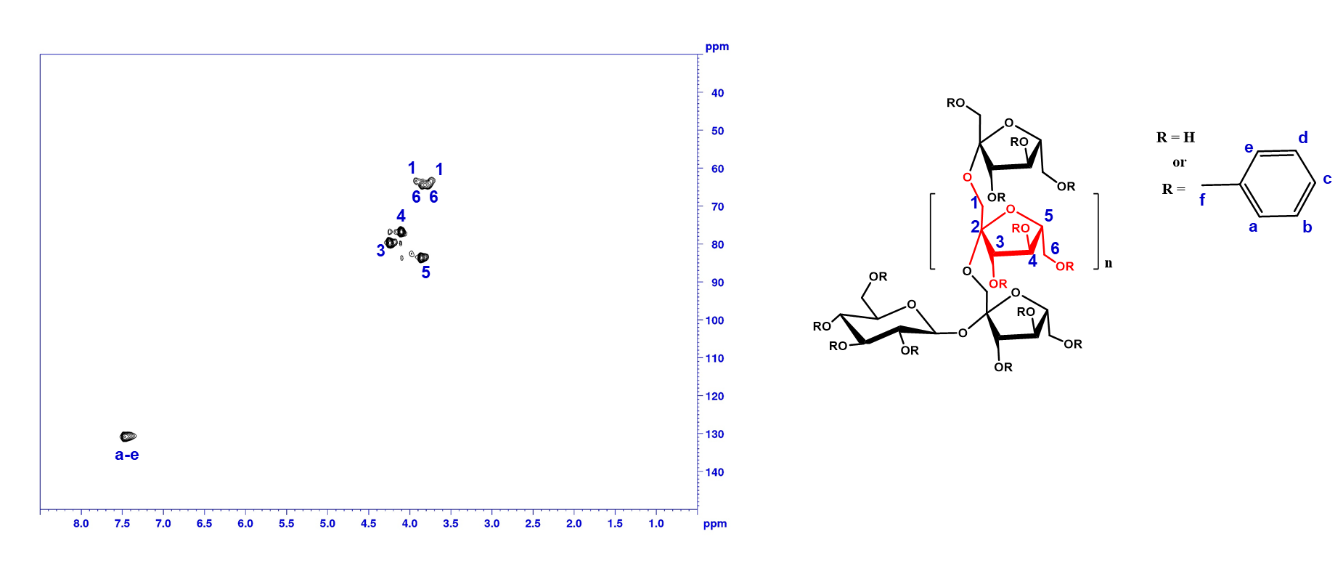

**Figure S6.** 2D ^13^C-^1^H HSQC NMR spectrum of benzyl-inulin (25 mg/ml) dissolved in D_2_O shown along with the resonance assignment. Peaks from methyl ^1^H-^13^C (labeled as ‘f’) from the R groups are not assigned due to spectral overlap.

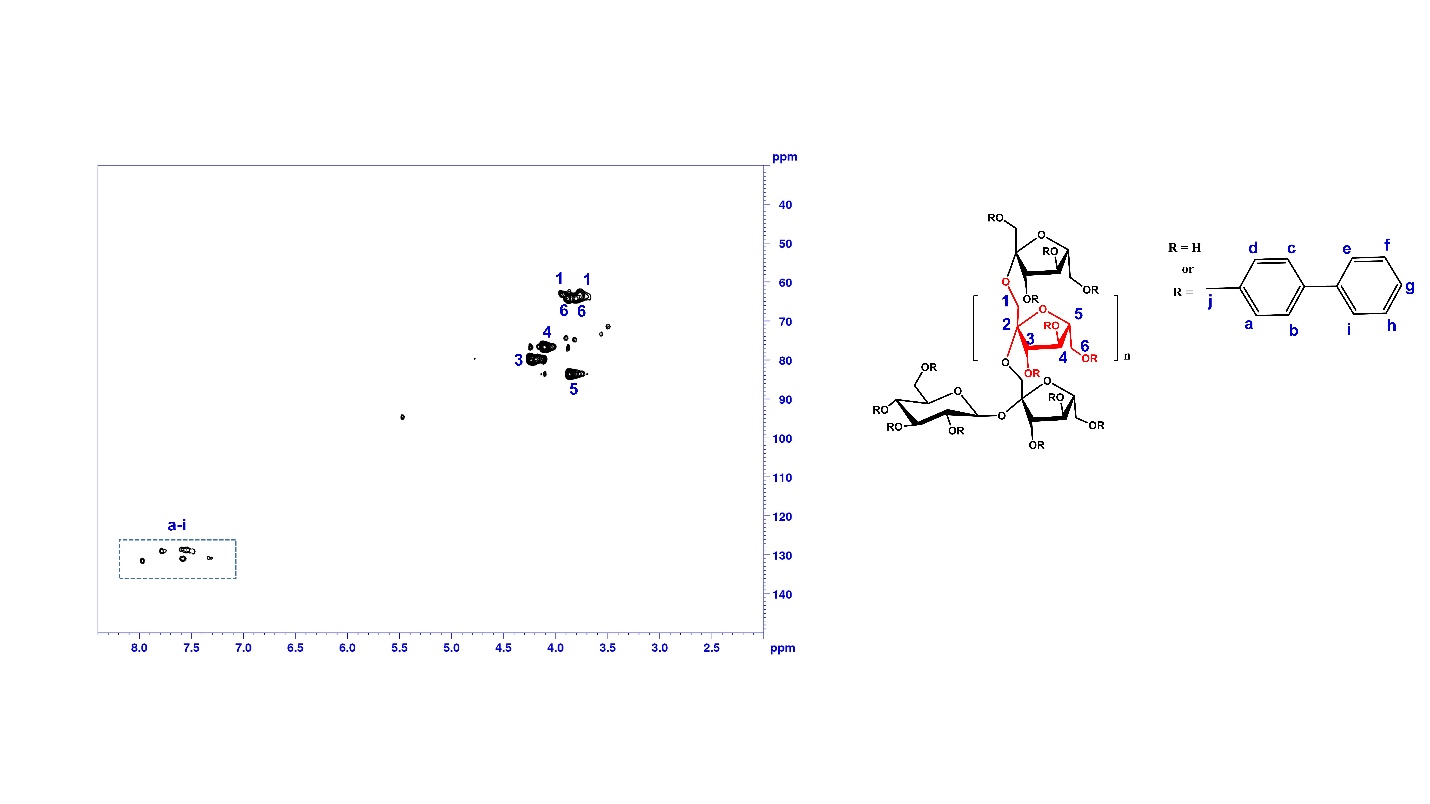

**Figure S7.** 2D ^13^C-^1^H HSQC NMR spectrum of biphenyl-inulin (25 mg/ml) dissolved in D_2_O shown along with the resonance assignment. Peaks from methyl ^1^H-^13^C (labeled as ‘j’) from the R groups are not assigned due to spectral overlap.

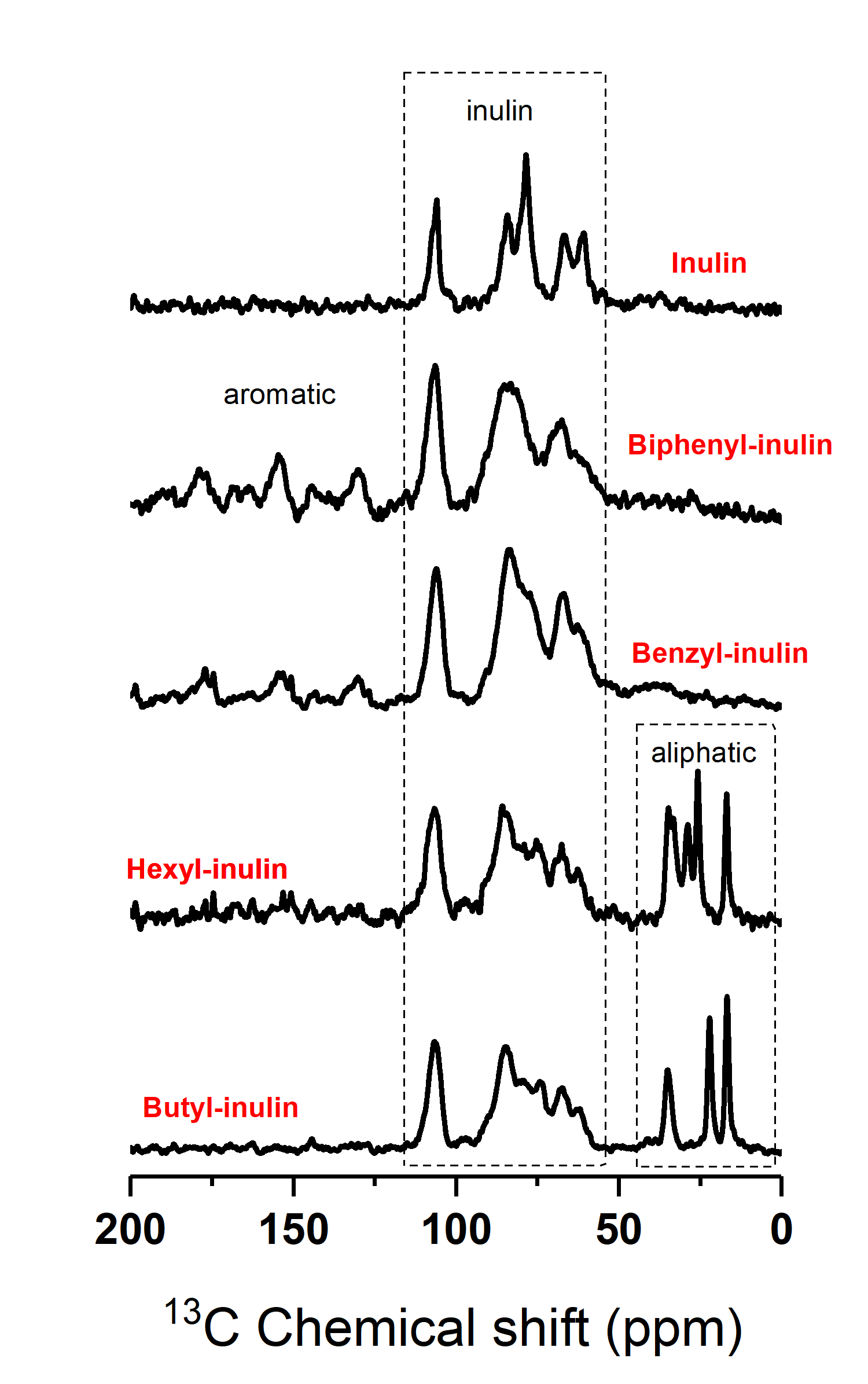

**Figure S8.** ^13^C CP-MAS NMR spectra of inulin and inulin derivatives acquired under 12 kHz MAS (magic angle spinning) showing the presence of peaks from the aliphatic and aromatic functional groups. These spectra were used to confirm the extent of functionalization of inulin.

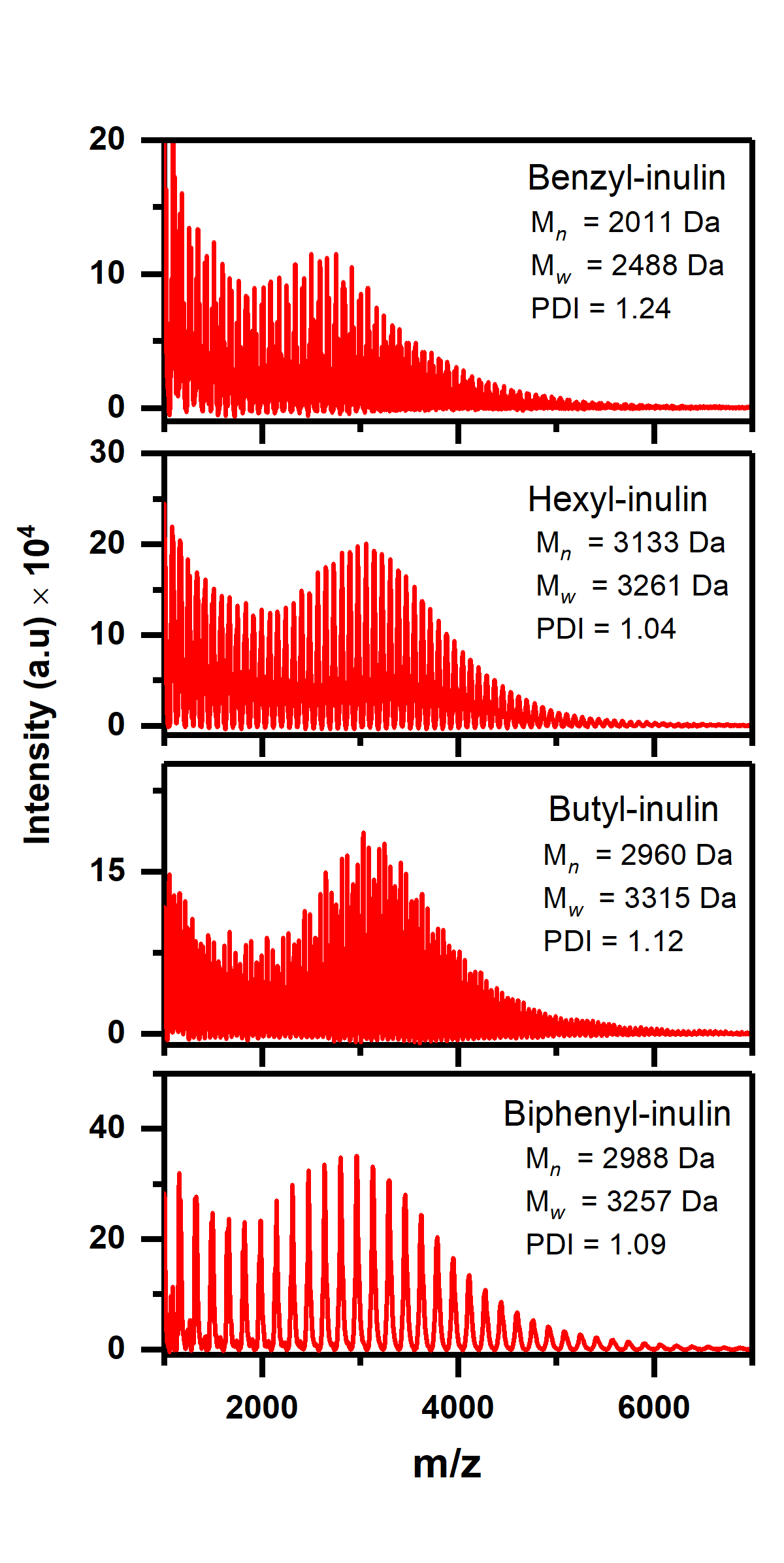

**Figure S9.** MALDI-MS spectra of inulin derivatives showing the polydispersity index (PDI) and the average molecular weight. Our estimation of the average molecular weight for inulin is comparable to the values reported in the literature.^[3-5]^

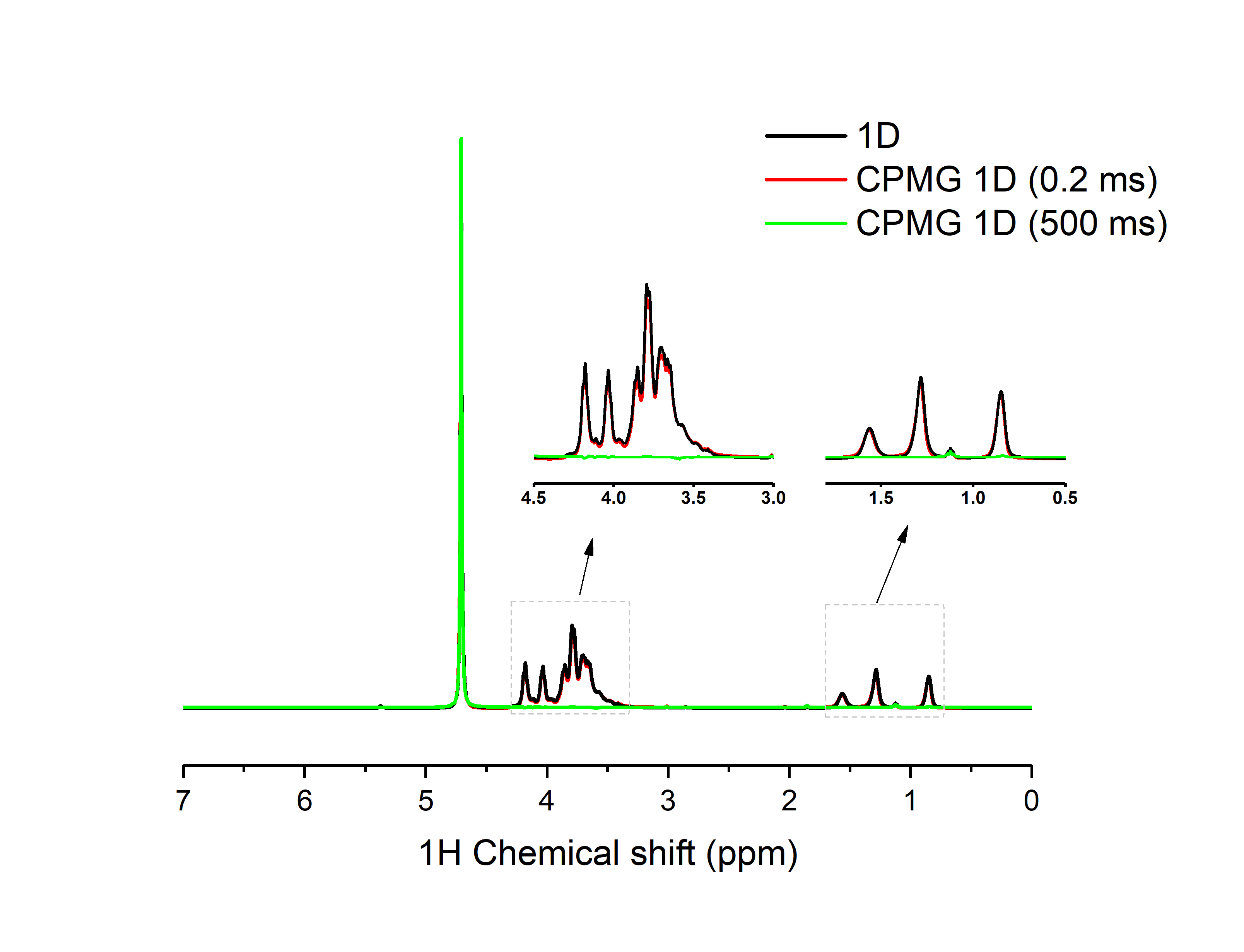

**Figure S10.** ^1^H NMR spectra of pentyl-inulin (10 mg/ml in D_2_O) without (black) and with the T_2_ relaxation filter (in CPMG (Carr-Purcell-Meibhoom-Gill)^[6]^ experiment) of 0.2 ms (red) and 500 ms (green) acquired at room temperature on a 500 MHz Bruker NMR spectrometer. All NMR spectra are normalized to the water-proton signal intensity. The T_2_ relaxation filter reduces the intensity of peaks from relatively fast relaxing components in the sample (or chemical groups in the molecule). While a short relaxation delay of 0.2 ms (red) did not affect the signal intensities, a long relaxation delay (i.e. 500 ms; the green NMR spectrum) completely suppressed the peaks from the polymer. In addition, the absence of any narrow peaks in the 500 ms relaxation delay spectrum (green) confirms the absence of any small molecular weight species in the solution.

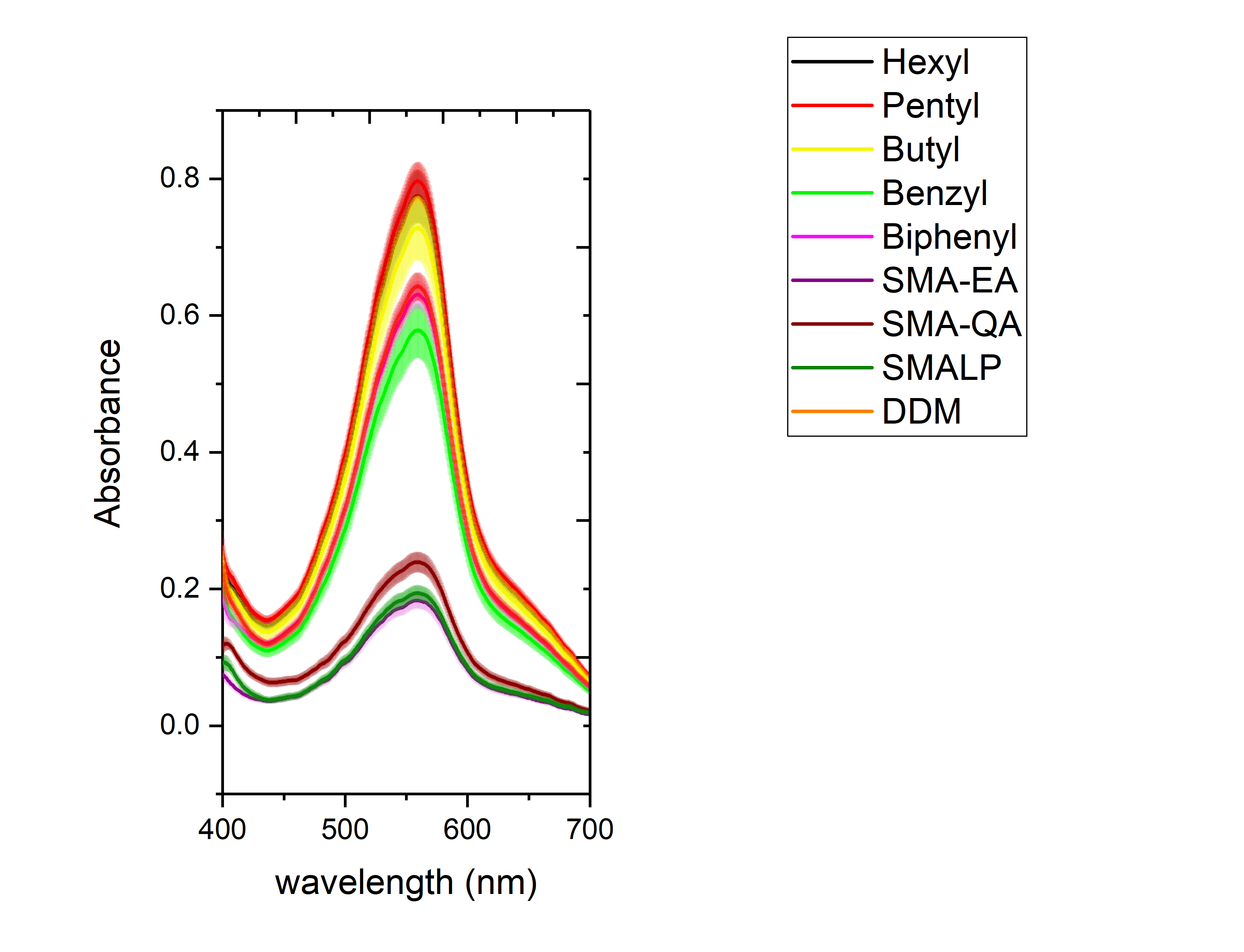

**Figure S11.** UV spectra obtained using Pierce BCA Protein Assay kit to quantify the efficiency of membrane protein solubilization by the synthetic polymers. The amount of solubilized protein content was obtained by measuring the absorption at 562 nm. The absorption values are normalized to 100% for DDM. See Figure 3 in the main text.
